# Biomechanical 3D tumor models on a micro-milled high-throughput force sensor array

**DOI:** 10.1101/2025.09.22.677940

**Authors:** Bashar Emon, Ahmadreza Kashefi, Md Habibur Rahman, Darbaz Adnan, Natalia Ospina-Munoz, Seamus Mellican, Sophia Santiago, William C Drennan, Md Saddam Hossain Joy, Aja A. Phan, Tasmia Afrin, Bumsoo Han, Kyle C Smith, Faraz Bishehsari, M Taher A Saif

**Affiliations:** Department of Mechanical Science and Engineering, Grainger College of Engineering, University of Illinois Urbana-Champaign, Urbana, IL 61801, USA; Rush Center for Integrated Microbiome and Chronobiology Research, Rush University Medical Center, Chicago, IL 60612, USA; Department of Health and Kinesiology, College of Applied Health Sciences, University of Illinois Urbana-Champaign, Urbana, IL 61801, USA; Department of Materials Science and Engineering, Grainger College of Engineering, University of Illinois Urbana-Champaign, Urbana, IL 61801, USA; Beckman Institute for Advanced Science and Technology, University of Illinois Urbana-Champaign, Urbana, IL 61801, USA; Computational Science and Engineering Program, Grainger College of Engineering, University of Illinois Urbana-Champaign, Urbana, IL 61801, USA; Gastroenterology Research Center, Division of Gastroenterology, Hepatology & Nutrition, Department of Internal Medicine, University of Texas Houston, TX 77030, USA; MD Anderson Cancer Center-UTHealth Houston Graduate School of Biomedical Sciences; CZ Biohub Chicago, LLC, Chicago, IL

**Author notes:** Equal contributing authors (Bashar Emon, Ahmadreza Kashefi, Md Habibur Rahman). Corresponding authors (Saif -; Bishehsari -; Smith -).

## Abstract

The tumor microenvironment plays a critical role in drug resistance, with extracellular matrix (ECM) mechanics, cell-cell crosstalk, and transport barriers contributing to poor therapeutic outcomes. Traditional two-dimensional (2D) cultures fail to capture these features, and drug efficacy in 2D often does not translate to three-dimensional (3D) models or in vivo tumors. Here, we introduce a 3D tumor model integrated with a high-throughput biomechanical sensor array that enables simultaneous measurement of cellular forces, matrix remodeling, and molecular transport. Fabricated using a scalable and cost-effective micro-milling approach, the platform allows parallel generation of multiple tumor constructs within a single dish. Using patient-derived pancreatic ductal adenocarcinoma (PDA) cells and stromal fibroblasts, we show that responses to gemcitabine and all-trans retinoic acid (ATRA) in 3D differ markedly from 2D cultures, consistent with clinical observations. By integrating biochemical and biomechanical readouts, this technology provides a more physiologically relevant tumor model and a powerful tool for preclinical drug testing and personalized medicine.

## INTRODUCTION

Traditional two-dimensional (2D) culture systems have been instrumental in cancer research and drug development, but they poorly recapitulate the complexity of *in vivo* tumor microenvironment [1]. In contrast, three-dimensional (3D) tissue models provide more physiologically relevant cell-cell and cell-matrix interactions, and therefore hold greater promise for drug screening and personalized medicine [2,3]. One major reason for this discrepancy is that 3D tissues preserve mechanical and structural features of the tumor microenvironment such as matrix stiffness, cell-generated forces, and dynamic cell-matrix crosstalk that profoundly affect drug penetration, signaling pathways, and therapeutic efficacy [4–6]. These biomechanical interactions, absent in 2D cultures, highlight the need for methods that can directly measure forces and mechanical remodeling in 3D models. However, while current 3D models capture biological and biochemical aspects, mechanobiological properties remain underutilized as facile and precise measurement of cellular forces within 3D extracellular matrices (ECM) have been a long-standing challenge [7–12]. Moreover, existing platforms that can measure such forces are often low-throughput and technically complex, limiting their scalability for systematic studies or clinical applications. Thus, there remains a critical need for accessible, high-throughput tools that integrate biomechanics into 3D disease models.

To address this, we introduced a 3D tumor model, integrated with a biomechanical sensor, that enables simultaneous probing of biochemical and biomechanical parameters, providing a comprehensive view of how cellular forces, matrix stiffness, and remodeling contribute to therapeutic response in 3D tumor models. The system is capable of quantifying: i) the force exerted by cells within 3D matrices and ii) the stiffness of the cell-matrix environment over time to assess matrix remodeling by cells [13,14]. This sensor circumvents the assumption of constitutive relations by using force balance to measure traction forces generated by small cell clusters in complex matrices. The sensors are made of polydimethylsiloxane (PDMS) cast from a microfabricated silicon mold [13,14]. Despite its successful implementation in research labs, the sensor’s relatively high cost and complexity have limited its widespread clinical adoption [15–17]. To enhance the sensor’s adaptability and ease of use, here we developed a high-throughput sensor array fabricated with PDMS using low-cost micro-milled polymethylmethacrylate (PMMA) molds. Compared to previous methods [7,13,18], this array of sensors facilitates generation of multiple replicates with a substantially faster workflow, enhancing versatility and enabling exploration of new avenues for applications beyond fundamental research, including potential integration into hospital drug screening protocols for individualized medicine by enabling the observation and quantification of critical biophysical features of the tumor microenvironment, including diffusivity, alongside the assessment of cellular biochemical signaling and viability, thereby providing valuable insights into its therapeutic relevance.

This innovative fabrication approach offers several advantages over other methods such as deep reactive ion etching (DRIE) or stereolithography [19]. Firstly, it allows the simultaneous generation of numerous samples within a single dish in a high throughput fashion, facilitating the exploration of diverse culture conditions. Secondly, micro-milling of PMMA significantly reduces production costs and time compared to traditional silicon microfabrication techniques like DRIE [19–21]. Thirdly, micro-milling enables greater depth of cutting and offers a larger space compared to etching silicon wafers with limited dimensions. Finally, milling techniques are capable of creating variable depth and three-dimensional structures without any additional steps. In contrast, DRIE requires multiple etching steps and masks to achieve similar complexity and depth, which can increase the time and cost of the fabrication process. The main limitation of micro-milling, compared to DRIE, is its precision and resolution). However, the precision of micro-milling is still better than that of 3D printing), making it a viable option for applications requiring moderate to high resolution. The high-throughput device, fabricated using micro-milled PMMA molds, combines cost-effectiveness, precision, and the ability to generate multiple replicates simultaneously. This approach significantly lowers production costs and enhances accessibility to advanced biomechanical tools. Despite the limitations in resolution, the benefits of reduced cost, space, depth of cut and three-dimensionality make micro-milling a promising method to fabricate precision equipment for testing biomaterials and cell/tissue mechanics. To demonstrate such capabilities, we utilized in vitro cancer tumors as a model system, since the dynamic interplay between cellular forces and the tumor microenvironment plays a critical role in cancer progression [22–26]. This approach allowed us to showcase the device’s ability to measure biomechanical parameters, such as contractility and matrix remodeling, providing valuable insights into cancer mechanobiology.

To demonstrate the utility of this platform beyond fabrication, we employed our high-throughput sensor array to probe the mechanobiology of 3D tumor tissues under drug treatment. Specifically, we tested the efficacy of all-trans retinoic acid (ATRA) and gemcitabine in patient-derived pancreatic ductal adenocarcinoma (PDA) organoids. The platform allowed us to monitor not only drug-induced changes in contractility and matrix remodeling over time, but also alterations in molecular diffusivity within the 3D constructs. These biomechanical and transport readouts provide an added dimension of insight compared to conventional viability-based assays, highlighting the importance of integrating mechanics into drug testing and discovery.

## RESULTS

### Concept: 3D tumor model assembled on a mechanical sensor

The unit sensor, housing a single *in vitro* tumor, is designed based on our previous work [13]. In concept, the sensor consists of three main components: a soft spring, a stiff spring, and two grips connected to the springs (Fig. 1A). The soft spring (*K*_*f*_) is the force sensor, and the stiff spring stabilizes the 3D tumor specimen that self-assembles in situ from a capillary bridge made with cell-ECM mixture between the grips (Fig. 1A). When cells generate contractile force *F*, it deforms the soft spring by *δ*_*f*_, thus producing a force readout from the sensor of *F*_*readout*_ = *K*_*f*_ ∗ *δ*_*f*_. In addition, by stretching or compressing the specimen, we can measure the mechanical properties (e.g., stiffness, elastic or visco-elastic moduli) of biomaterials and matrices. We designed the springs as cantilever frames made of PDMS (Fig. 1A). The spring frame, consisting of thin beams, is designed and anchored at one end so that the other end (connected to the specimen) can translate along the length of the tissue sample. The spring constant for n beams is *K*_*s*_ = ^12*nEI*^⁄_*L*3_; where E is the modulus of elasticity of PDMS, *I* is the moment of inertia (*I* =ℎ*b*^3^ ⁄_12_, with b and h being the width and depth of the beams), and *L* is the length of the beams.

**Fig. 1:**
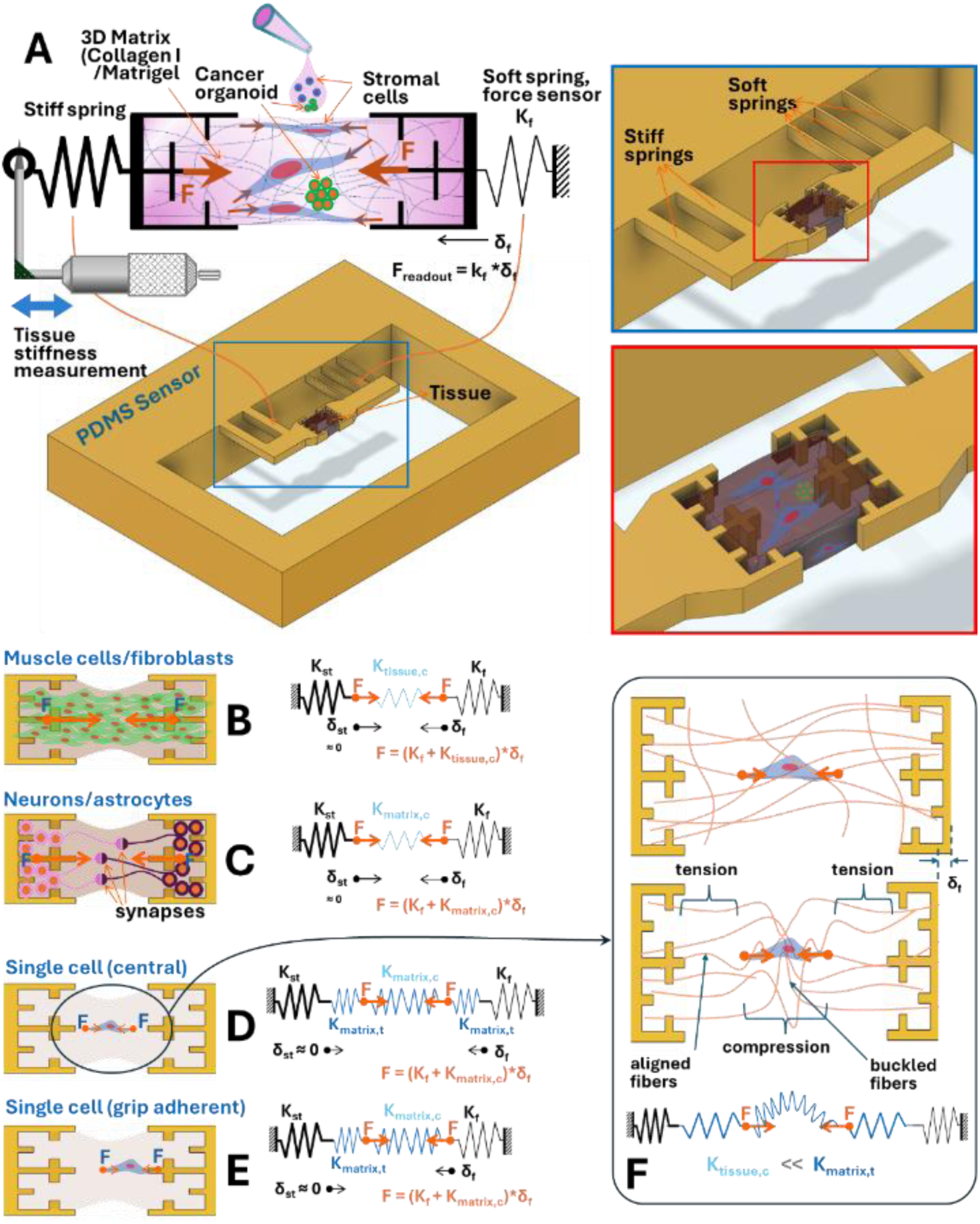
Concept and design of a sensor for measuring cell generated force in 3D matrices and tissue stiffness. (A) Schematic illustration of specimen formation and working principles of the sensor. The tissue sample self-assembles on the sensor from a capillary bridge of cell-ECM mixture between the grips. Over time, the cells activate, interact with the ECM fibers, and generate contractile forces, which are transferred to and deform the soft spring (with a spring constant of *K_f_*). Spring deformation and cellular activity are monitored using a microscope. Cell force is determined by multiplying the spring constant by the deformation *(K_f_*δ_f_)*. Tissue stiffness is measured by applying axial compression or tension on the specimen with a micrometer attached to a stiff beam, while spring deformation and grip movement are continuously tracked to measure force and strain. For the PDMS sensor, soft springs are designed as thin beams, and wide beams represent stiff springs. Force transmission mechanism in tissues with (B) high density muscle cells or fibroblasts, (C) neurons connected to both grips, (D) single cell adherent to one grip, and (E-F) single cell located at the center. For cases in B-C, *F = (K_f_ + K_tissue,c_)*δ_f_ ≈ K_f_ *δ_f_* as *K_tissue,c <<_ K_f_*. For cases in C-D, *F = (K_f_ + K_matrix,c_)*δ_f_ ≈ K_f_ *δ_f_* as *K_matrix,c <<_ K_f_*. Therefore, the force readout from the sensor, *F_readout_ = K_f_ * δ_f_* is a precise estimate of cell-generated force.

In order to understand the relationship between the force applied by the cell(s) within the matrix (*F*) and the force detected by the sensor (*F*_*readout*_), it is important to consider how the applied force is transmitted to the sensor spring. For example, a specimen with muscle cells forms bundles of myotubes that are attached to the grips on both ends (Fig. 1B). Such muscle tissues apply contractile force directly to the sensor springs through the grips and the total force generated by the muscle is *F* = (*K*_*f*_ + *K*_*tissue*,*c*_) ∗ *δ*_*f*_, where *K*_*tissue*,*c*_ is the stiffness of the tissue in compression. Typically, *K*_*tissue*,*c*_ ≪ *K*_*f*_, to produce an applied force *F* ≈ *K*_*f*_ ∗ *δ*_*f*_ = *F*_*readout*_. Similarly, for neuronal tissues (Fig. 1C), where cells are confined within the grips and neurites connect in the central region, force is applied directly on the force sensing spring which precisely detects the applied force. Here, *F* = (*K*_*f*_ + *K*_*matrix*,*c*_) ∗ *δ*_*f*_, where *K*_*matrix*,*c*_ is the stiffness of the matrix in compression. Again, when *K*_*matrix*,*c*_ ≪ *K*_*f*_ is satisfied, *F* ≈ *F*_*readout*_ is produced.

In specimens with single cells (Fig. 1D-E), the cells apply force on the matrix that transmits the force to the springs. When the cell is located at the center (Fig. 1E), the matrix within the force dipole is under compression. Matrices such as collagen are composed of thin fibers (∼ 0.1–1 μm [27]) that are strong in tension but buckle under small compressive stress. Therefore, any force on collagen is transferred almost entirely to the sensor’s spring by tension (Fig. 1F). It should be noted that a small fraction of the applied force is used to buckle collagen fibers, and this fraction is not transferred to the sensor spring, even for cases where the cell is adhering to the grips at one end (e.g. Fig. 1D) [14].

The sensor also enables mechanical testing of the specimens allowing for the measurement of mechanical properties such as elastic modulus or stiffness [13,14]. As illustrated in Fig. 1A, the method for assessing tissue stiffness involves stretching or compressing the tissue sample by moving the stiff spring with a 3D linear stage equipped with micrometers or piezo actuators. The change in length due to compression or tension, along with the corresponding force readout from the soft spring, provides information to determine the tissue’s stiffness. For a linear force-displacement response, tissue stiffness can be calculated directly. In cases where the relationship is non-linear, the force-deformation curve is fitted to an appropriate non-linear model to determine the tissue stiffness.

### Micro-milling for the fabrication of sensors with intricate structures

Leveraging recent progress in precision micro-milling technology [19], we investigated the feasibility of fabricating micron-scale structures and components that comprise biomechanical sensors. We fabricated PMMA molds for the components of the high-throughput sensor system (Fig. 2A). We employed two different methods to make the PMMA molds-direct and indirect milling. Direct milling uses end-mills with diameters smaller than the smallest feature in the design, creates grooves in the form of sensors and the device parts are cast directly from the PMMA mold. On the other hand, indirect milling can utilize larger end-mills to create an inverted model of the design (i.e. ridges in the shape of sensors) by cutting out the surrounding material. A secondary mold is then cast with PDMS from the PMMA model and then silanized with chlorotrimethylsilane to prevent adhesion with PDMS cured in contact. These molds are used to cast PDMS into arrays of sensors.

**Fig. 2:**
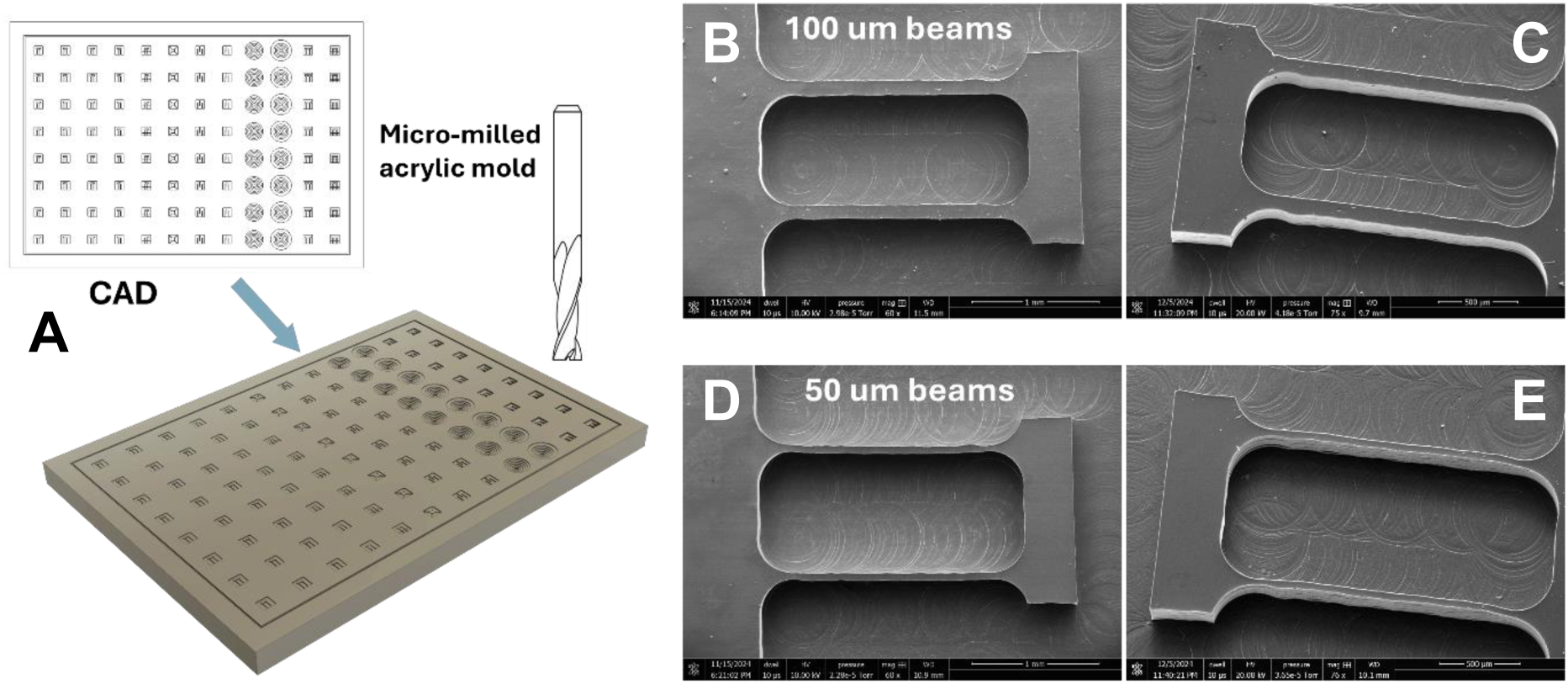
Capabilities of the present micro-milling technique in creating structures and components for biomechanical sensors. (A) An array of devices can be set up in the dimensions of a 96-well plate and fabricated from a micro-milled mold. Scanning electron microscope images of (B,C) 100 µm beams and (D,E) 50 µm beams fabricated from micro-milled PMMA mold.

The performance of the fabricated sensor components using these micro-milling techniques exhibits distinct characteristics. Direct milling often results in wall slopes and achieving a beam width smaller than 80 µm is challenging since smaller end-mills easily break while cutting hard PMMA. Additionally, the maximum aspect ratio (height/width of beams) attainable with direct milling is approximately two, which limits the height of the features relative to their width.

In contrast, with indirect milling, it is possible to achieve a beam width as small as 50 µm, and a remarkable aspect ratio of up to fifteen, allowing for the creation of much taller features relative to their width. Moreover, the sidewalls produced through indirect milling are nearly vertical, contributing to more precise and defined cross-sections (Fig. 2B,C). Another advantage of indirect milling is that the cutting volume is smaller when fabricating molds of sensor arrays. As a result, it takes less time to complete, making it a more efficient process. Furthermore, molds created using this technique can be utilized to make many replicates of the sensor array in PDMS, since PMMA resists wear for repeated use. Therefore, Tool wear and machining time are no longer significant concerns with this approach, as each mold only needs to be machined once.

However, a key limitation is that the gap between beams cannot be made smaller than the size of the tool bit used, which may restrict the design possibilities for certain applications. These differences in performance make indirect milling a more suitable choice for applications requiring high aspect ratios, finer resolution, and faster fabrication times, whereas direct milling is more appropriate for simpler designs with lower aspect ratio requirements.

### Assembly and operational setup

To make a functional high-throughput system from the micro-milled molds, we fabricated an array of precision sensors (Fig. 3A) and the top/bottom layers that create independent wells for each sensor. The sensor layer is sandwiched between the top and bottom layers and attached to a glass slide to construct the functional high-throughput device (Fig. 3B-D). The components are joined with PDMS to ensure a proper seal that prevents leakage and contamination between the wells.

**Fig. 3:**
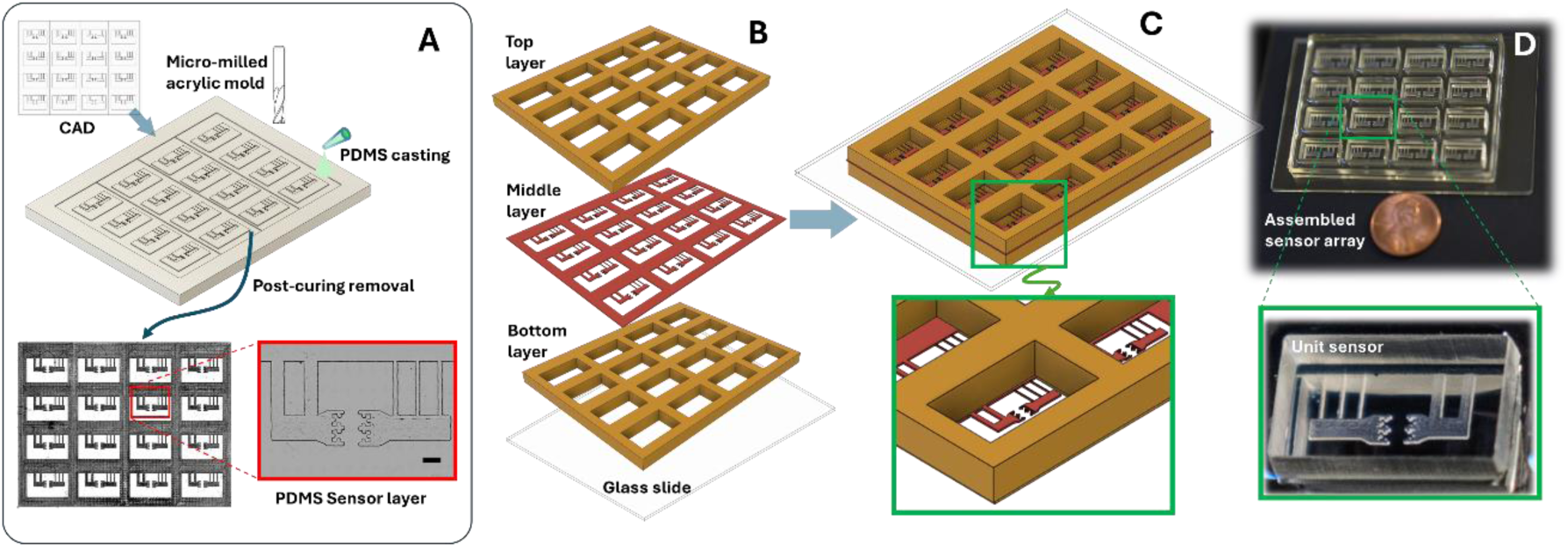
Fabrication and assembly of a high-throughput sensor array. (A) Molds for the device components are made by micro-milling PMMA sheets using computer-aided design and manufacturing. Liquid PDMS is carefully cast into the PMMA molds, cured, and peeled to make three separate components – top, bottom, and the middle sensor layers. Brightfield images of the sensor layer underscore the precision of fabrication. (B-C) Assembling the different layers on a glass slide creates the device with separate wells containing unit sensors. (D) Photographs of a functional device with a four-by-four sensor array.

Fig. 4 illustrates the process and outcomes of forming tissues on a sensor array. To enhance the sensor’s attachment with the matrices (e.g. collagen), the surfaces are treated with (3-aminopropyl)triethoxysilane (APTES) and glutaraldehyde, which improve hydrophilicity and enhance the bond between the grips and the ECM (Fig. 4A). This surface treatment also facilitates the flow of ECM into the micro-channels on the grips improving mechanical anchoring. Tissues are created by forming capillary bridges between the grips using a cell-ECM mixture, which self-assembles into structured tissues upon polymerization (Fig. 4B). Brightfield microscopy verified the successful formation of sixteen tissue samples, comprising collagen I and PSCs, distributed across a 4×4 sensor array (Fig. 4C). Subsequent Second Harmonic Generation (SHG) imaging provided insights into collagen organization within the tissues (Fig. 4D). The analysis revealed that PSCs significantly remodel collagen, as evidenced by increased SHG signal intensity (Fig. 4E) and a pronounced alignment of collagen fibers across the grips (Fig. 4F), underscoring their role in ECM remodeling.

**Fig. 4:**
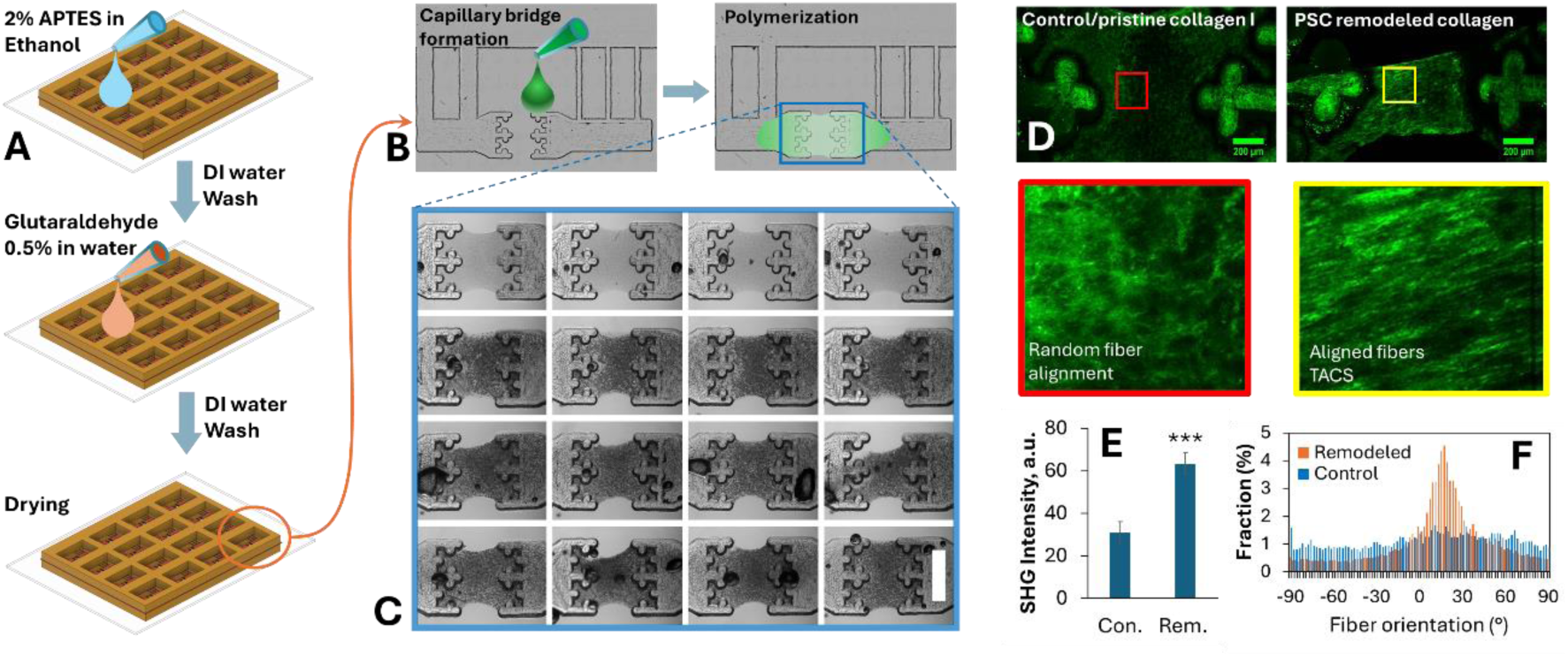
Forming tissues on the sensor array. (A) Sensors are treated with 2% APTES (v/v in ethanol) for 12 hours and then 0.5% glutaraldehyde (v/v in water) for an hour, to make the surfaces hydrophilic and to strengthen grip-ECM bonds. (B) Tissue specimens are formed by creating capillary bridges between the grips with cell-ECM mixture that self-assembles into a tissue upon polymerization. (C) Brightfield images of 16 tissue samples (collagen I + pancreatic stellate cells (PSCs) on a single device with 4×4 sensors. (D) Second Harmonic Generation (SHG) images of collagen in an acellular control sample and a PSC tissue. (E) SHG intensity and (F) fiber orientation histogram reveal that PSCs remodel collagen by compaction and fiber alignment.

### Research and clinical applications

#### Drug-response in 2D

The mechanical properties of the ECM support cancer development and contribute to treatment failure and immune evasion, making it a significant target for cancer therapies. Traditional 2D culture models fail to capture these critical mechanical and structural cues, and numerous studies have shown that drug efficacy in 2D often does not translate to 3D models or in vivo tumors [28–32]. In contrast, patient-based in vitro models, such as 3D tumor spheroids, have advanced our ability to predict drug response. However, better mimics of the complex interactions between the ECM and primary cancer cells are still highly needed. Consistent with previous findings [28–32], our results on PDA models show that therapeutic responses differ substantially between 2D and 3D, underscoring the relevance of biomechanical context in drug evaluation (Fig. 5).

**Fig. 5:**
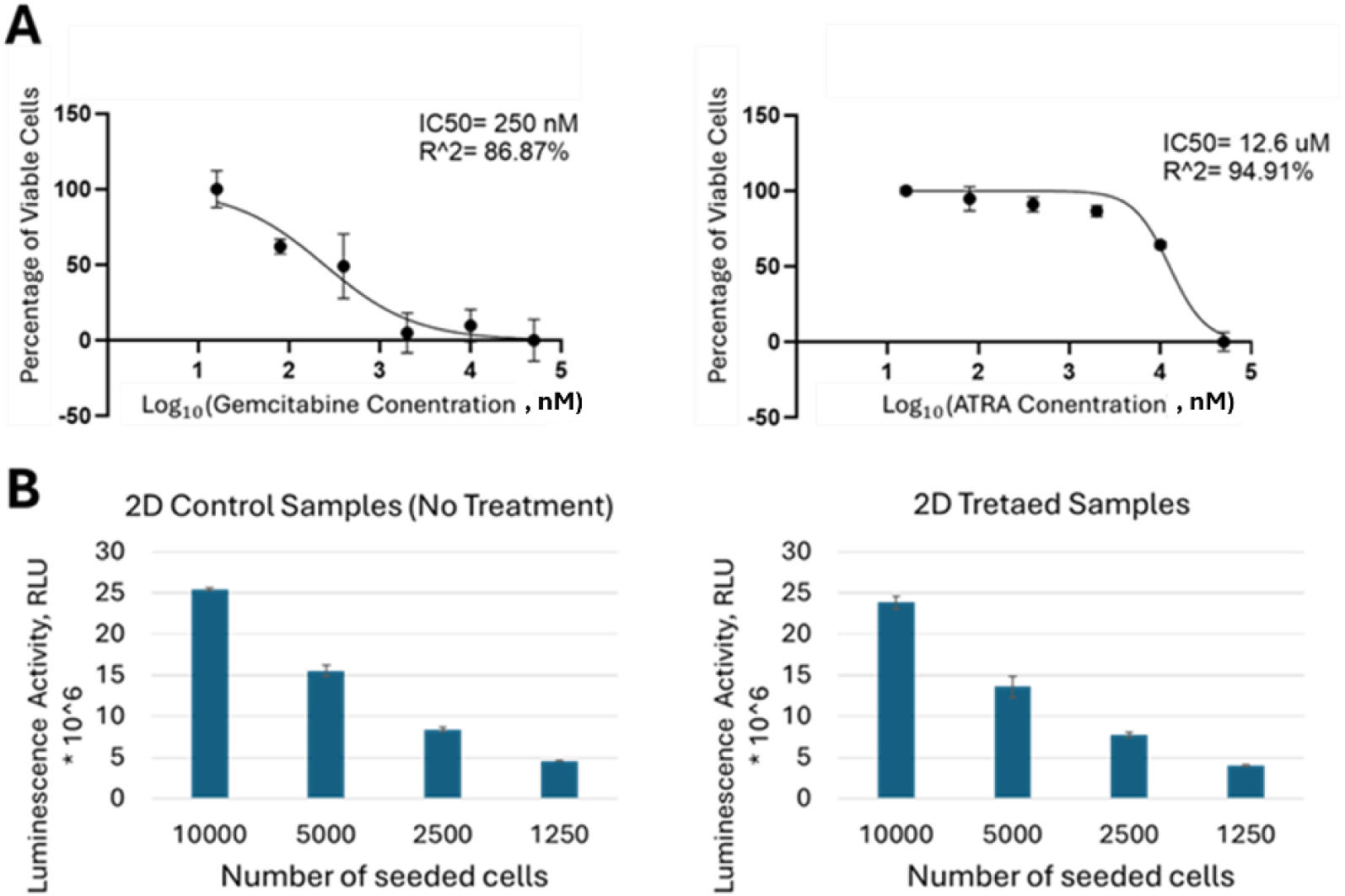
Dose-response of cells in pancreatic cancer microenvironment in 2D. (A) Monoculture of MIA PaCa II cancer cells treated with Gemcitabine exhibited an IC₅₀ of 250 nM, while pancreatic stellate cells (PSCs) treated with ATRA exhibited an IC₅₀ of 12.6 μM. (B) In 2D co-culture systems, treatment with Gemcitabine + ATRA at their IC₅₀ concentrations resulted in a about 10% reduction in luminescence signal across different seeding densities. Total number of cells shown in the figure, 2D co-cultures consisting of MIA PaCa II and PSCs in 2:1 ratio. Statistical significance was determined by student’s T-test (* p<0.1, ** p<0.01). Column plots: mean ± SD.

To establish the appropriate drug concentrations for assessing cell viability in both 2D and 3D culture systems, we performed half-maximal inhibitory concentration (IC₅₀) assays to determine the cytotoxicity profiles of the drugs. Dose-response analysis with 2D cultures indicated an IC₅₀ of 250 nM for Gemcitabine (a standard chemotherapeutic agent for pancreatic cancer [33–35]) in reducing MIA PaCa II cell activity by 50% (Fig. 5A). In addition, all-trans retinoic acid (ATRA) was examined due to its reported capacity to suppress PSC proliferation and activity and its emerging role as an investigational agent targeting the desmoplastic stroma [36–39]. We determined its IC₅₀ as 12.6 μM in inhibiting PSC activity by 50% in 2D PSC-only culture (Fig. 5A).

However, in 2D co-culture of MIA PaCa II cells and PSCs (2:1 ratio), treatment with Gemcitabine + ATRA at their IC₅₀ concentrations resulted in about 10% reduction in viability (Fig. 5B, which drug in 5B?), across different total cell number present in the co-culture, indicating decreased drug efficacy in the presence of PSCs.

#### Drug and Biomechanical response in 3D

To assess whether this effect was maintained in 3D cultures, micro-tumor models (co-culture of MIA PaCa II and PSCs (2:1 ratio)) were created on the microarray devices and allowed to compact for 24 hours prior to treatment. They were then exposed for 24 hours to Gemcitabine, ATRA, or Gemcitabine + ATRA at their respective IC₅₀ concentrations, with untreated tissues as controls (Fig 6A). Viability was measured using the CellTiter-Glo 3D assay, after 24 hours of tissues being treated with all of the drug regimens (Fig. 6F). Utilizing our high-throughput sensor system with cancer models *in vitro* (Fig. 6), we investigated the biomechanical and drug response of the 3D cancer tissue consisted of MIA PaCa II and PSCs. To establish a model that recapitulates key features of the pancreatic cancer microenvironment, we developed a co-culture system consisting of pancreatic cancer cells (MIA PaCa II) and PSCs with 2:1 ratio (Fig. 6A-E). The cells were allowed to compact for 24 hours prior to any treatment. In this setting, cancer cells act as the primary drivers of PSC activation, while PSCs contribute to ECM remodeling, a critical process in pancreatic tumor progression.

**Fig. 6:**
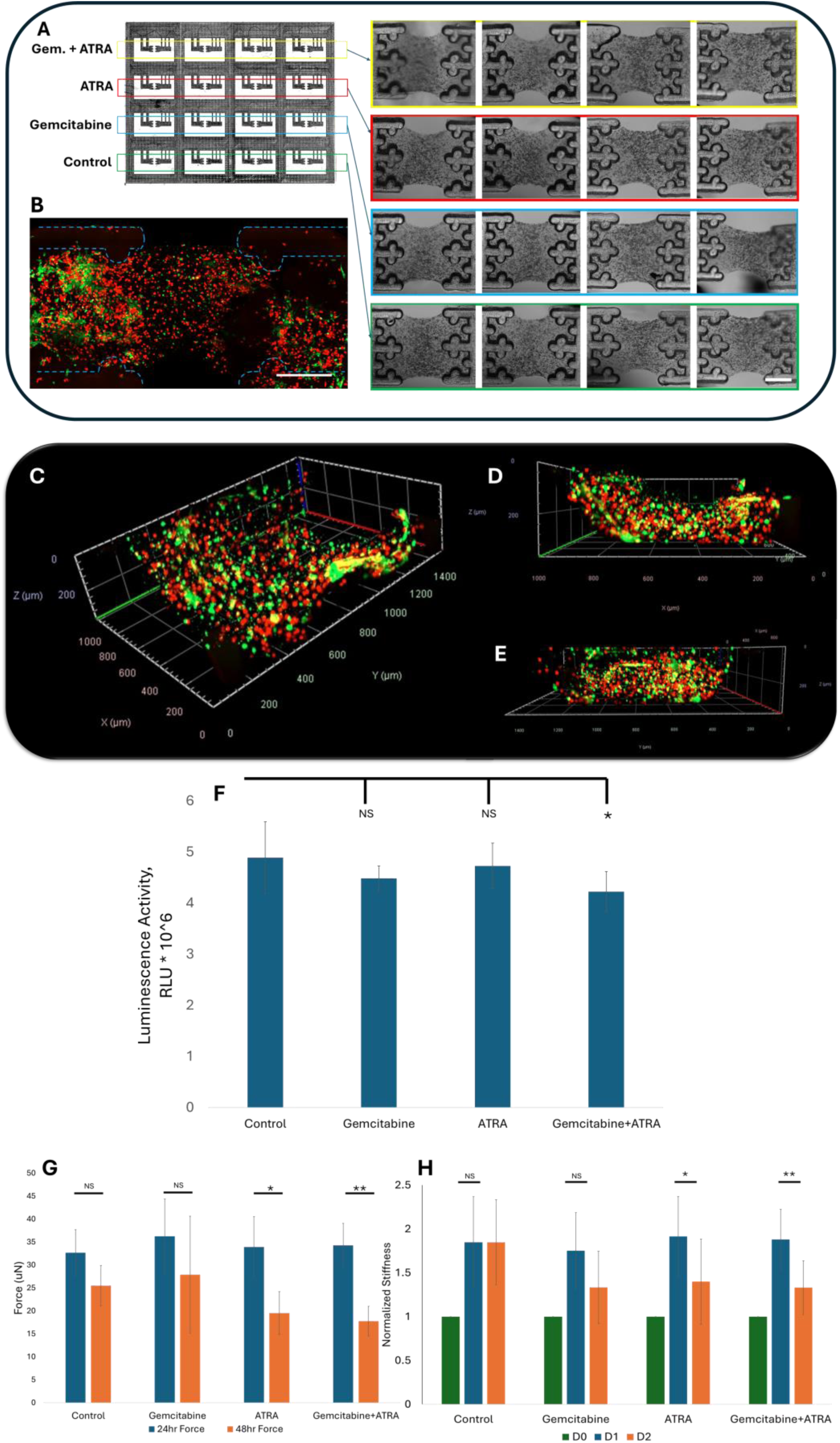
Configuration of 3D co-culture tissues consisting of a co-culture of MIA PaCa II and PSCs with a concentration of 1.32 M/ml and 0.66 M/ml, respectively. (A) Experimental setup for drug administration after the initial compaction of the tissues and brightfield (BF) images of tissue specimens containing a co-culture of MIA PaCa II and in 2 mg/ml of Collagen Type I and 1 mg/ml Matrigel, before compaction. (B) Fluorescence microscopy image of a tissue, with PSCs marked with Green CellTracker and MIA PaCa II cells secreting tdTomato fluorescent protein. 3D fluorescence images of the tissue construct between the grips. The tissue was fixedafter two hours of the tissue fomation, and fluorescence confocal microscopy was performed directly after fixation. The grip area has been subtracted from the image. (C) Isometric view of the tissue, with tdTomato-expressing MIA PaCa II cells (red) and PSCs labeled with Green Cell Tracker (green). (D) Orthogonal view showing the 3D distribution of cells along the X-axis. (E) Orthogonal view showing the 3D distribution of cells along the Y-axis. (F) Viability of the cells in 3D tissue shows no significant effect with Gemcitbaine or ATRA monotherapy but a significant decline in viability with the treatment of Gemcitabine+ATRA, with all drug treatments being performed at their IC₅₀ concentrations. (G) Administration of ATRA, and its combination with Gemcitabine decrease tumor force significantly. 24hr Force (blue bar) and 48hr Force (orange bar) indicate the force generated by the tissues after 24 hours of compaction and after 24 hours of being treated with the drugs, respectively. (H) Matrix remodeling by the co-culture can be reversed by the application of ATRA and a combination of Gemcitabine and ATRA. D0 (green bar), D1 (blue bar), and D2 (orange bar) are normalized stiffness of the matrix right after being constructed, after 24 hours of compaction (without any drug administration), and after 24 hours of drug administration (all drugs and drug combination at their IC50 value), respectively. Statistical significance was determined by student’s T-test (* p<0.05, ** p<0.005). Column plots: mean ± SD. Scale bar: 500 *μ*m.

The tissues were treated and exposed to Gemcitabine, ATRA, or Gemcitabine + ATRA at their respective IC₅₀ concentrations for the next 24 hours, with untreated tissues as controls (Fig 6A). Viability was measured using the CellTiter-Glo 3D assay. There was no statistically significant change in viability due to ATRA, or Gemcitabine alone. Gemcitabine + ATRA reduced viability by about 13.5%.

Within 24 hours, PSC-mediated contractile activity generated significant force (Fig. 6F) and increased scaffold stiffness by approximately 2-fold, indicating ECM remodeling (Fig. 6DG).

Drug treatment for 24 hours revealed distinct mechanobiological outcomes. Treatment with ATRA, either alone or in combination with Gemcitabine, significantly decreased tumor force (Fig. 6F) and administration of a combination of Gemcitabine and ATRA attenuated matrix stiffness significantly. By contrast, Gemcitabine monotherapy,failed to induce any significant change in either force or stiffness. These findings suggest that Gemcitabine-ATRA-mediated stromal reprogramming exerts a pronounced influence on the properties of the pancreatic cancer microenvironment, an effect not replicated by Gemcitabine alone.

#### Patient-derived 3D PDA organoids for biomechanical response to chemotherapy

To demonstrate the translational capabilities of the system, we utilized patient-derived organoids (PDOs) to model pancreatic adenocarcinoma on the high-throughput sensor array (Fig. 7A). We chose PDOs because these exhibit active stroma, with some tumors exhibiting mechanical stiffness up to ten times greater than normal tissue [22,24,26], and hold tremendous promise for individualized medicine in cancer [40,41]. To scale up the tumor model, we co-cultured PDOs with human pancreatic stellate cells (PSCs), which are key cells contributing to stroma formation in the pancreas. Inclusion of PSCs into our PDA model increased mechanical forces significantly (Fig. 7B). Stiff stroma is thought to negatively impact the drug efficacy in PDA. We hypothesized that targeting PSCs with ATRA can augment the effect of gemcitabine, a standard chemotherapy in PDA, by reducing the forces. Interestingly, greater reduction in contractility and stiffness was observed with time when Gemcitabine (2 µM) was combined with ATRA (3.3 µM) Fig. 7B-C, Table 1). These results suggest that the high-throughput system is capable of clinical applications such as personalized medicine.

**Fig. 7:**
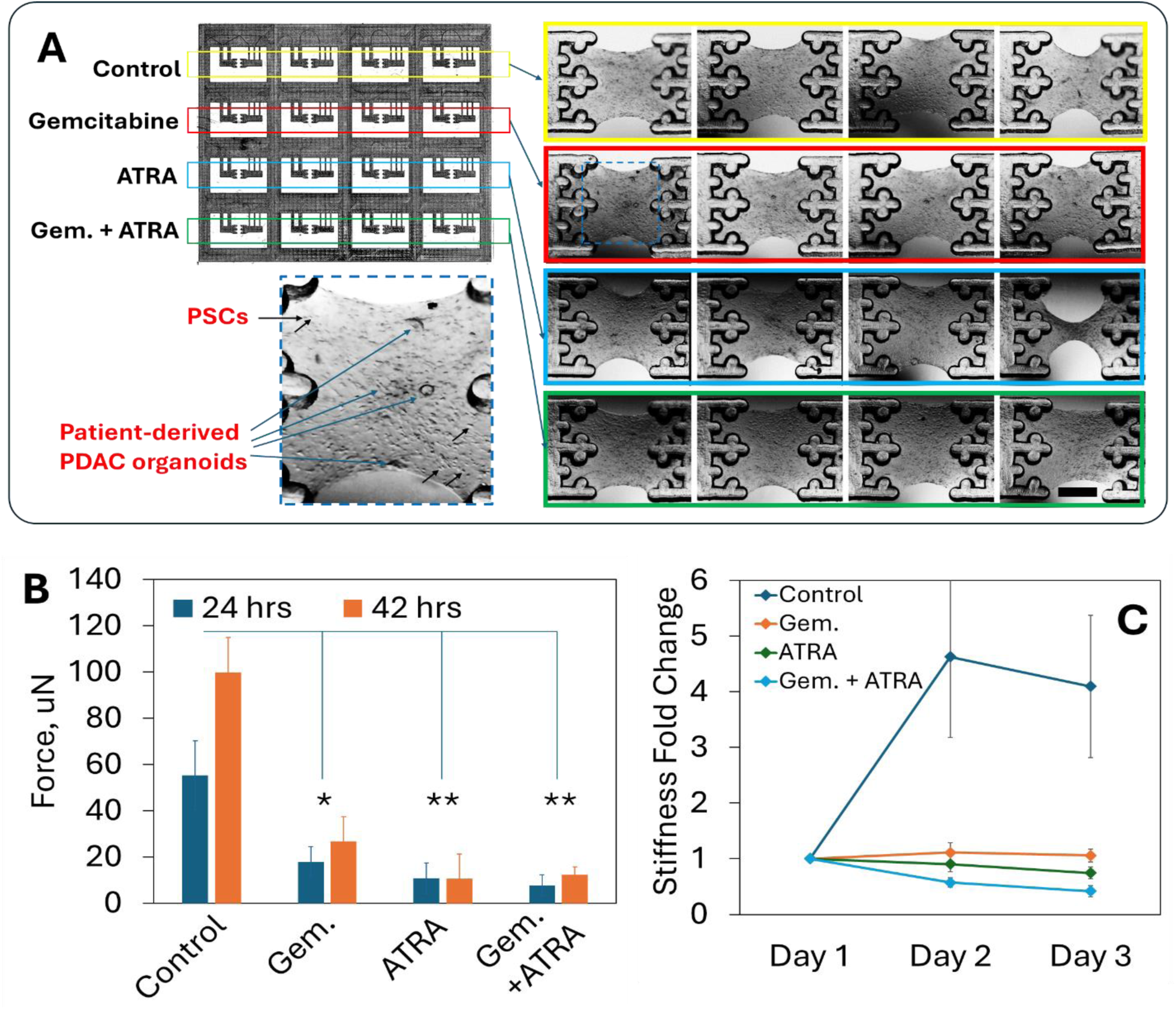
Effects of chemotherapy drugs on patient-derived PDAC models. (A) Experimental setup for drug administration and brightfield images of *in vitro* PDAC tissues with patient-derived organoids and PSCs. (Inset) Some of the organoids (blue arrows) and PSCs (black arrows) are highlighted. (B) Comparison of tissue force at 24 and 42 hours. (C) Change of tissue stiffness in these patient-specific tumor models. Stiffness of the tissues were normalized with respect to Day 1 stiffness of the same sample. Gemcitabine and ATRA both inhibit matrix stiffening and contractility; however, combination treatment is more efficient. Error bars: SD. Statistical significance was determined by student’s T-test (* p<0.05, ** p<0.01). Scale bar: 500 μm.

**Table 1:**
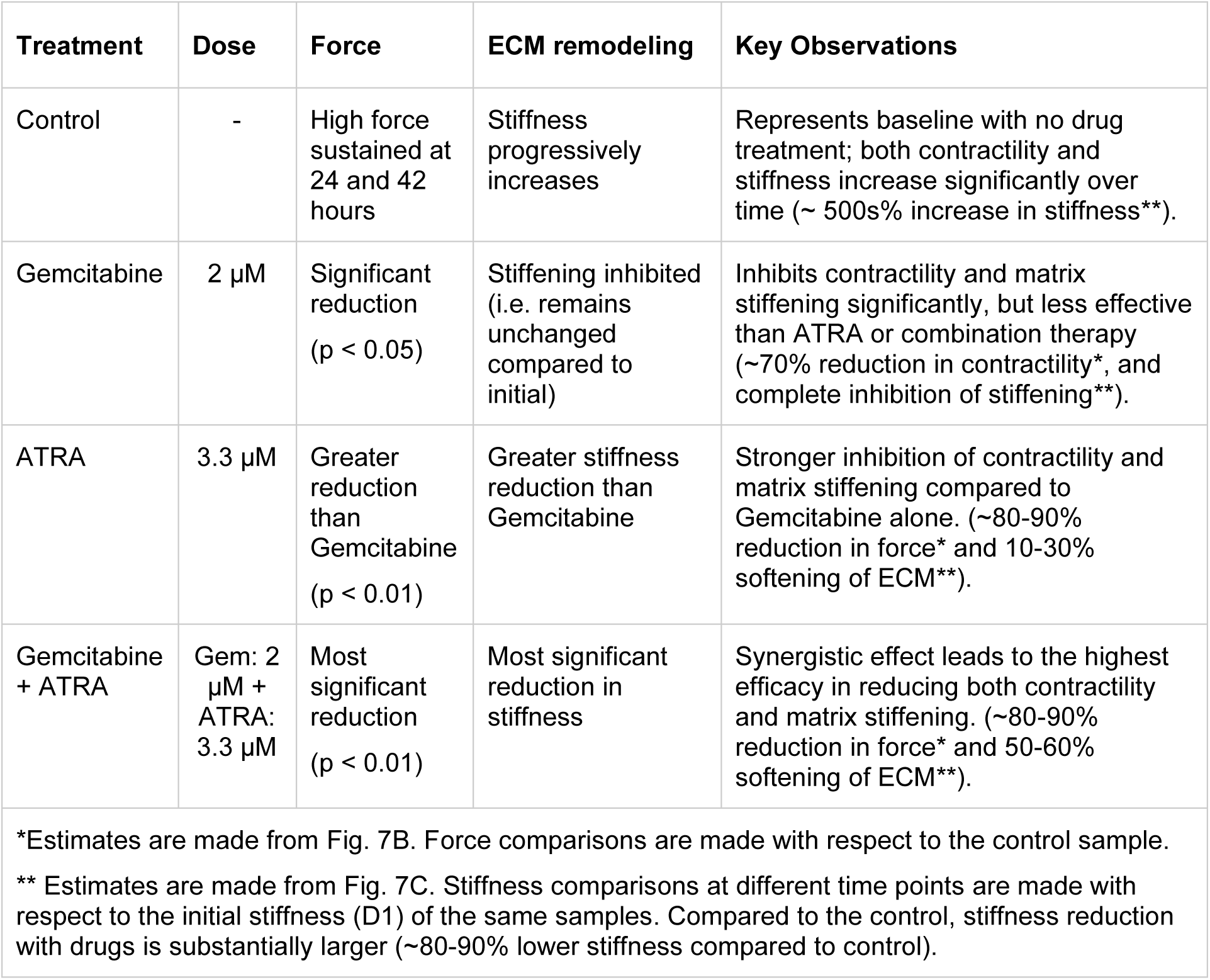
Biomechanical outputs of drug treatment of *in vitro* PDA models.

### Performance comparison and future directions

Despite the significant advancements presented by our high-throughput sensor array, some limitations suggest scope for improvement. While micro-milling offers a cost-effective and scalable alternative to traditional microfabrication methods, our current technology lacks the ultra-high precision (∼1 µm), particularly when compared to lithographic techniques like DRIE that can achieve feature sizes down to the sub-micron level and aspect ratio of ∼ 20:1 [42,43]. In contrast, using micro-milling, we created 50(±5) micron features with a 15:1 aspect ratio. 3D printers, capable of producing large objects (e.g. 128mm x 85mm for a 96-well plate), can achieve the smallest feature size of ∼ 200 microns with a maximum aspect ratio of ∼ 20 [44]. Therefore, micro-milling can fill the gap in resolution between lithography and 3D printing and is significantly cost-effective [19–21]. In addition, it enables deeper structures (depth >1 mm), that are not feasible with DRIE. Moreover, creating 3D features with micro-milling is considerably faster (one-step process) than with lithography, which requires multiple steps adding to time and cost.

Another limitation stems from the constraints associated with using smaller-sized endmills in the micro-milling process. As the diameter of the endmills decreases to achieve finer features, there is an increased susceptibility to tool wear and breakage, which can compromise the dimensional accuracy and consistency of the fabricated sensors. Additionally, smaller endmills require longer machining times, which can lead to thermal effects that potentially distort the PMMA molds. Hence, it is important to use endmills suitable for the instrument and material. Also, the system should have a mechanism for chip removal and temperature control.

There are several avenues for future research and development. One direction is to refine the micro-milling process to enhance the precision of the sensor components, potentially through the integration of advanced milling techniques or hybrid fabrication approaches that combine micro-milling with other high-resolution methods. Another promising direction is the development of more complex tissue models on the sensor array, incorporating multiple cell types, ECM components, and dynamic biochemical gradients to better mimic *in vivo* conditions. This could involve the integration of vascular networks or immune cell populations, providing a more comprehensive understanding of how different cell types interact within a three-dimensional matrix under various mechanical and biochemical cues.

Furthermore, expanding the application of the sensor array beyond basic research to include more clinically relevant scenarios, such as patient-specific drug screening or real-time monitoring of tissue-engineered constructs, could significantly enhance its utility. Collaborations with clinical researchers and industry partners could facilitate the translation of this technology into routine clinical practice, potentially revolutionizing the way we assess tissue mechanics and drug efficacy in personalized medicine.

Moreover, this system enables more comprehensive investigations of the tumor microenvironment. By incorporating key cell types involved in cancer progression, it facilitates fundamental studies of the underlying mechanisms of tumor development. The findings presented here highlight the potential of this platform for probing biophysical cues relevant to cancer research. In particular, the system offers a direct means to examine how cells and the extracellular matrix (ECM) interact and collectively drive biophysical alterations within the tissue.

Therefore, while our high-throughput sensor array represents a major step forward in the field of mechanobiology, ongoing improvements in fabrication techniques, model complexity, and clinical integration will be crucial to fully realize its potential in both research and clinical applications.

## DISCUSSION

The high-throughput sensor array provides versatile and scalable technology to probe mechanobiology, particularly in the context of healthy or diseased tissues reliant on cellular forces and matrix mechanics. This innovation addresses long-standing challenges in quantifying the forces exerted by cells or tissue constructs in vitro, as well as assessing the dynamic remodeling of the cell-matrix environment. By leveraging micro-milling techniques to fabricate low-cost, scalable, and versatile PDMS molds, we have created a system that not only facilitates high-throughput experiments but also holds the potential for integration into clinical settings, such as drug screening protocols. The sensor array’s ability to generate multiple 3D samples simultaneously within a single dish enhances the experimental throughput, while the cost-effectiveness and time efficiency of micro-milling make it an attractive alternative to more expensive and time-consuming lithographic techniques. Additionally, the capacity to create deeper, three-dimensional structures with variable depths, without requiring multiple processing steps, provides a level of versatility that is particularly valuable for complex tumor modeling.

However, there are limitations inherent to the current micro-milling technology that must be addressed. The precision and resolution achievable with micro-milling, while slightly superior to 3D printing in this case, do not yet match the ultra-high precision offered by DRIE, especially for applications requiring micrometer-scale features. The challenges associated with using smaller-sized endmills, including tool wear, breakage, and thermal distortion of PMMA molds, further highlight the need for ongoing refinement of the fabrication process. Future research should focus on optimizing these processes, possibly through the integration of hybrid fabrication techniques that combine the strengths of micro-milling with other high-resolution methods.

In terms of impact, the potential applications of this sensor array extend far beyond fundamental research. Our results in the successful demonstration of its use with patient-derived cancer organoids underscore its utility in faithfully modeling tumor-stroma interactions in 3D and in individualized drug efficacy testing. This was exemplified in studies with ATRA, which effectively reverses PSC-induced matrix stiffening by reducing contractility and enhancing proteolytic remodeling, thereby normalizing the tumor microenvironment [45]. These findings open the possibility of reducing matrix stiffness with ATRA to enhance tissue permeability and amplify the efficacy of chemotherapy when used in combination. While promising, this hypothesis warrants further investigation to validate its therapeutic potential. The ability to assess mechanobiological responses in real time within 3D tumor models presents a powerful tool for personalized medicine, offering insights that could guide therapeutic decision-making. Consequently, our high-throughput sensor array represents a promising advance for both research and clinical applications in the emerging field of mechanobiology.

## MATERIALS AND METHODS

### Cell culture

Human pancreatic stellate cells (PSCs, ScienCell Cat# 3830) were cultured in Stellate Cell Medium (SteCM, Cat #5301) as previously described by Haque et al [40]. Cells were grown at 37 °C in a humidified incubator with 5% CO_2_.

MIA PaCa II cells were grown in a high-glucose Dulbecco’s Modified Eagle’s Medium (DMEM) (Corning, Cat#: 10-013) supplemented with 10% FBS (Gibco, Cat#: A5670701) and 1% Penicilin/Streptomycin (Gibco, Cat#: 15140122).

### Fabrication and assembly of PDMS sensors

The sensors were cast from micro-milled PMMA molds. These molds were designed in Fusion 360, and G-code was generated for milling them from clear PMMA. 6 mm thick scratch- and UV-resistant PMMA sheets (McMaster-Carr) were machined with a micro-end mill (Harvey Tool, USA) with a cutter diameter and length of cut of about 0.508 mm and 1.524 mm, respectively. A desktop computer numerical control (CNC) machine (Bulkman 3D Ultimate Bee) equipped with precision ball screws and driven by four closed-loop stepper motor using GRBL Mega v1.1 motion control firmware [46] was employed for this task. 24,000 rev/min spindle speed was used along with a maximum feed rate of 45.72 cm/min for positioning after manual collet adjustment to minimize tool runout. The dimensions of the machined cross-sections were measured using ImageJ software.

PDMS (Sylgard 184) was prepared by mixing the base and cross-linker at a 10:1 ratio by weight. Next, PDMS was pipetted into the molds, allowed to fill all the features and trenches by capillary micro-molding [47], and then cured at 60 °C for 24 hours. Using sharp tweezers, these device components were carefully peeled off the molds. For assembling the setup, the three layers of the device were joined by uncured PDMS, ensuring minimal air entrapment and a proper seal. Next, the assembled part was autoclaved, treated with 2% (v/v in ethanol) APTES (Sigma-Aldrich) for 12 hours (and three times wash in ethanol after) and 0.5% (w/v in water) glutaraldehyde (Electron Microscopy Sciences) for 1 hour (and 3x wash in water after). This part was then dried and glued to a large microscope slide (Ted Pella) using uncured PDMS. The device was ready to use after the PDMS glue was cured at 60 °C.

### Tissue formation

High-concentration rat-tail collagen I (Corning, Cat#: 354249) solution was prepared on ice by neutralizing 9.5 mg/ml stock with 1N sodium hydroxide, 10X PBS, and deionized (DI) water with proportions recommended in Corning protocol [48] to achieve a concentration of 4 mg/ml ECM with pH 7.2. The tissue precursor solution was prepared by mixing cell suspension (in media) with collagen at a 1:1 ratio to achieve a final collagen concentration of 2 mg/ml and cell density of 150 x10^3^ cells/ml. This density results in <10 cells in the tissue. To produce a higher number of cells, we increased density linearly based on the desired number of cells in the final tissue construct. Cell-ECM mixture was then pipetted onto the grips of the sensors, and a capillary bridge was formed. With time, the solution filled the channels in the grips and polymerized at 37 °C for 45 minutes. It is important to add a small volume of media to the corners of each well before forming tissues, to prevent drying of tissues. After polymerization, culture media was carefully added to the wells with sensors so that the assembled tissues were inundated and then the array was placed in the incubator or microscope for experimentation.

Matrigel (Corning, Cat#: 354237) was incorporated in selected experiments. In these cases, the cell suspension in culture medium was combined with Matrigel to achieve a concentration of 2 mg/ml. The resulting cell-Matrigel suspension was subsequently mixed with collagen at a 1:1 ratio, resulting in a final concentration of 1 mg/ml for Matrigel.

### Chemicals and Drugs

Gemcitabine (MedChemExpress, Cat #: HY-17026) was reconstituted in Dimethyl Sulfoxide (DMSO) to a stock concentration of 10 mM and stored at −80 °C until use. All-Trans Retinoic Acid (ATRA) (Sigma, Cat#: R2625) was prepared similarly in DMSO at 10 mM and maintained at −80°C until experimental application.

### IC50 Determination, Cell Viability, and Microplate Reading

The half-maximal inhibitory concentration (IC₅₀) of gemcitabine and ATRA was determined using the alamarBlue HS Cell Viability Reagent (Invitrogen, Cat#: A50100). Cells were seeded with an initial number of 5000 cells per well in an opaque-walled microplate in triplicates. All of the cells were allowed to adhere to the bottom of the microplate for the first 24 hours. Cells were then treated with a range of drug concentration from 50 uM to 16 nM with 5-fold change between each drug concentration. Cells were then incubated and treated with the drugs for 48 hours. Cells were then treated with a volumetric ratio of 2:1 (reagent: medium), followed by fluorescence measurement using the GloMax Explorer Microplate Reader (Promega Corporation).

Cell viability in 2D and 3D cultures was assessed using the CellTiter-Glo 3D Cell Viability Assay (Promega, Cat#: G9681).

2D cultures were seeded with a total number of 5000 cells per well consisting of both MIA PaCa II and PSCs in a 2:1 ratio in triplicates. Cells were then allowed to adhere to the bottom of the opaque-walled microplate for 24 hours. After the initial 24 hours 2D co-cultures were treated and then incubated with Gemcitabine + ATRA at their IC_50_ concentrations for 48 hours. For viability measurement, 2D cultures were incubated with the reagent at a 2:1 ratio (reagent: medium) and shaken for 30 minutes. Luminescence, as a surrogate for viability, was then recorded on the GloMax Explorer (Promega Corporation).

For quantifying the viability of cells in 3D, tissues were formed on the micro-array devices, resulting in 8 samples per each drug regimen, using two micro-array devices in parallel. The tissues were allowed to compact and remodel the matrix for the first 24 hours. The tissues were then treated with Gemcitabine, ATRA, and Gemcitabine+ATRA, all at their IC_50_ concentrations, with 8 samples being kept as controls and not treated with any drugs.For viability measurement, 3D tissues were incubated with the reagent at a 2:1 ratio (reagent: medium) and shaken for 30 minutes. The supernatant was then transferred to an opaque-walled plate, and luminescence was recorded on the GloMax Explorer (Promega Corporation).

### Patient-derived pancreatic ductal adenocarcinoma organoids

Patient-derived organoid (PDO) was generated as described in previous studies [40,49]. Organoids were generated from biopsies of untreated pancreatic cancer tissue at Rush University Medical Center. In brief, the biopsy tissues were digested using collagenase II, Dispase II, and DNase I, then washed multiple times. They were subsequently plated in 50 μL Matrigel domes on a 24-well plate with 500 μL of organoid medium. The growth medium consisted of Advanced DMEM/F12, N2 Supplement, N-acetyl cysteine, B27, PGE2, HEPES, nicotinamide, gastrin, hEGF, A83-01, Y-27632, hFGF, and Wnt3A–R-spondin 1–Noggin conditioned media (50% of the final volume).

### Imaging

The experiment was run in an environment-controlled chamber enclosing an inverted optical microscope (Olympus IX81, 10X objective, Olympus) mounted on a vibration isolation table (Newport Corporation). The chamber maintains cell culture conditions at 37 °C temperature, 5% CO_2_ and 70% humidity. Moreover, a motorized stage (Prior Scientific Inc.) was programmed to track multiple specimens at preset locations at multiple time points. Images of both the tissues were acquired in brightfield mode with a Neo sCMOS camera (Andor Technology). For calculating spring displacements, images of the sensor gauges were analyzed using a template matching plugin in ImageJ with sub-pixel resolution. Force and stiffness calculations were made from the spring displacements, according to a previously described protocol [14].

SHG images were acquired subsequently using a LSM 710 (Zeiss) two-photon excitation microscope and 20X objective lens. Maximum intensity projection of the acquired confocal z-stacks was constructed using the ImageJ (U.S. National Institutes of Health) software.

Fluorescence microscopy of the 3D cancer tissue constructs was performed using a LSM 980 confocal microscope (Zeiss) equipped with a 20X objective lens.

## ACKNOWLEDGEMENTS

Research reported in this publication was partially supported by NSF grant ECCS 1934991, Cancer Center at Illinois (CCIL) seed grant, and the Chan Zuckerberg Biohub Chicago. F.B. would like to acknowledge NIH support, CA27948, CA277110 and AG086145. K.C.S. and M.H.R. were supported by the U.S. Office of Naval Research (Award No. N00014-22-1-2577).

## AUTHOR CONTRIBUTIONS

B.E., M.T.A.S. and F.B. conceived and designed the experiments. B.E., M.H.R., W.C.D. and K.C.S. fabricated the sensors. B.E., A.K., A.A.P. and D.A. performed the experiments. B.E., A.K., A.A.P., M.S.H.J. performed imaging and analysis. N.O. and S.M. helped with viability experiments. S.S. fabricated the micro-array devices. T.A. summarized the drug testing results in a tabular form. B.E., A.K., F.B. and M.T.A.S. prepared the manuscript. All authors have read, edited and approved the final manuscript.

## DATA AVAILABILITY

All data needed to evaluate the conclusions in the paper are present in the paper and/or the Supplementary Materials. The authors will make available any additional data/information related to the paper upon request.

## SUPPLEMENTARY

## REFERENCES

[1] G. Rauner, P.B. Gupta, C. Kuperwasser, From 2D to 3D and beyond: the evolution and impact of in vitro tumor models in cancer research, Nat Methods (2025) 1–12. 10.1038/S41592-025-02769-1;SUBJMETA.

[2] W.H. Abuwatfa, W.G. Pitt, G.A. Husseini, Scaffold-based 3D cell culture models in cancer research, J Biomed Sci 31 (2024) 1–39. 10.1186/S12929-024-00994-Y/TABLES/8.

[3] Z. Peng, X. Lv, H. Sun, L. Zhao, S. Huang, 3D tumor cultures for drug resistance and screening development in clinical applications, Mol Cancer 24 (2025) 1–16. 10.1186/S12943-025-02281-2/TABLES/1.

[4] M. Sheth, M. Sharma, M. Lehn, H. Al Reza, T. Takebe, V. Takiar, T. Wise-Draper, L. Esfandiari, Three-dimensional matrix stiffness modulates mechanosensitive and phenotypic alterations in oral squamous cell carcinoma spheroids, APL Bioeng 8 (2024) 36106. 10.1063/5.0210134.

[5] A. Micalet, E. Moeendarbary, U. Cheema, 3D In Vitro Models for Investigating the Role of Stiffness in Cancer Invasion, ACS Biomater Sci Eng 9 (2021) 3729–3741. 10.1021/ACSBIOMATERIALS.0C01530.

[6] M.H. Zaman, L.M. Trapani, A. Siemeski, D. MacKellar, H. Gong, R.D. Kamm, A. Wells, D.A. Lauffenburger, P. Matsudaira, Migration of tumor cells in 3D matrices is governed by matrix stiffness along with cell-matrix adhesion and proteolysis, PNAS July 18 (2006) 10889–10894. www.pnas.orgcgidoi10.1073pnas.0604460103 (accessed September 14, 2025).

[7] B.A. Nerger, M.J. Siedlik, C.M. Nelson, Microfabricated tissues for investigating traction forces involved in cell migration and tissue morphogenesis., Cell Mol Life Sci 74 (2017) 1819–1834. 10.1007/s00018-016-2439-z.

[8] J.C. del Álamo, R. Meili, B. Álvarez-González, B. Alonso-Latorre, E. Bastounis, R. Firtel, J.C. Lasheras, Three-Dimensional Quantification of Cellular Traction Forces and Mechanosensing of Thin Substrata by Fourier Traction Force Microscopy, PLoS One 8 (2013) e69850. 10.1371/JOURNAL.PONE.0069850.

[9] A. Hayn, T. Fischer, C.T. Mierke, Inhomogeneities in 3D Collagen Matrices Impact Matrix Mechanics and Cancer Cell Migration, Front Cell Dev Biol 8 (2020) 593879. 10.3389/FCELL.2020.593879/BIBTEX.

[10] R.J. Polackwich, D. Koch, R. Arevalo, A.M. Miermont, K.J. Jee, J. Lazar, J. Urbach, S.C. Mueller, R.G. McAllister, A Novel 3D Fibril Force Assay Implicates Src in Tumor Cell Force Generation in Collagen Networks, PLoS One 8 (2013) e58138. 10.1371/JOURNAL.PONE.0058138.

[11] D. Böhringer, M. Cóndor, L. Bischof, T. Czerwinski, N. Gampl, P.A. Ngo, A. Bauer, C. Voskens, R. López-Posadas, K. Franze, S. Budday, C. Mark, B. Fabry, R. Gerum, Dynamic traction force measurements of migrating immune cells in 3D biopolymer matrices, Nature Physics 2024 20:11 20 (2024) 1816–1823. 10.1038/s41567-024-02632-8.

[12] J. Steinwachs, C. Metzner, K. Skodzek, N. Lang, I. Thievessen, C. Mark, S. Münster, K.E. Aifantis, B. Fabry, Three-dimensional force microscopy of cells in biopolymer networks, Nat Methods 13 (2016) 171–176. 10.1038/nmeth.3685.

[13] B. Emon, Z. Li, M.S.H. Joy, U. Doha, F. Kosari, M.T.A. Saif, A novel method for sensor-based quantification of single/multicellular force dynamics and stiffening in 3D matrices, Sci Adv 7 (2021) eabf2629. 10.1126/sciadv.abf2629.

[14] B. Emon, M.S.H. Joy, W.C. Drennan, M.T.A. Saif, A multi-functional sensor for cell traction force, matrix remodeling and biomechanical assays in self-assembled 3D tissues in vitro, Nature Protocols (in Press) (2024).

[15] M.M. Eikanger, K.S. Schraufnagel, J.L. Slunecka, R.A. Potts, J. Freeling, R.L. Brockstein, B. Emon, M.T.A. Saif, K. Rezvani, Veratridine, a plant-derived alkaloid, suppresses the hyperactive Rictor-mTORC2 pathway: a new targeted therapy for primary and metastatic colorectal cancer, (n.d.). 10.21203/rs.3.rs-5199838/v1.

[16] B. Emon, M.S.H. Joy, L. Lalonde, A. Ghrayeb, U. Doha, L. Ladehoff, R. Brockstein, C. Saengow, R.H. Ewoldt, M.T.A. Saif, Nuclear deformation regulates YAP dynamics in cancer associated fibroblasts, Acta Biomater 173 (2024) 93–108. 10.1016/J.ACTBIO.2023.11.015.

[17] M. Saddam Hossain Joy, D.L. Nall, B. Emon, K. Yun Lee, A. Barishman, M. Ahmed, S. Rahman, P.R. Selvin, M.T.A. Saif, Synapses without tension fail to fire in an in vitro network of hippocampal neurons, Proceedings of the National Academy of Sciences 120 (2023) e2311995120. 10.1073/PNAS.2311995120.

[18] W.R. Legant, A. Pathak, M.T. Yang, V.S. Deshpande, R.M. McMeeking, C.S. Chen, Microfabricated tissue gauges to measure and manipulate forces from 3D microtissues, Proceedings of the National Academy of Sciences 106 (2009) 10097–10102. 10.1073/pnas.0900174106.

[19] D.J. Guckenberger, T.E. De Groot, A.M.D. Wan, D.J. Beebe, E.W.K. Young, Micromilling: a method for ultra-rapid prototyping of plastic microfluidic devices, Lab Chip 15 (2015) 2364–2378. 10.1039/C5LC00234F.

[20] D. Serje, J. Pacheco, E. Diez, Micromilling research: current trends and future prospects, The International Journal of Advanced Manufacturing Technology 2020 111:7 111 (2020) 1889–1916. 10.1007/S00170-020-06205-W.

[21] B.Z. Balázs, N. Geier, M. Takács, J.P. Davim, A review on micro-milling: recent advances and future trends, The International Journal of Advanced Manufacturing Technology 2020 112:3 112 (2020) 655–684. 10.1007/S00170-020-06445-W.

[22] B. Emon, J. Bauer, Y. Jain, B. Jung, T. Saif, Biophysics of Tumor Microenvironment and Cancer Metastasis - A Mini Review, Comput Struct Biotechnol J 16 (2018) 279–287. 10.1016/j.csbj.2018.07.003.

[23] J. Bauer, M.A.B. Emon, J.J. Staudacher, A.L. Thomas, J. Zessner-Spitzenberg, G. Mancinelli, N. Krett, M.T. Saif, B. Jung, Increased stiffness of the tumor microenvironment in colon cancer stimulates cancer associated fibroblast-mediated prometastatic activin A signaling, Sci Rep 10 (2020) 1–11. 10.1038/s41598-019-55687-6.

[24] A.J. Rice, E. Cortes, D. Lachowski, B.C.H. Cheung, S.A. Karim, J.P. Morton, A. del Río Hernández, Matrix stiffness induces epithelial–mesenchymal transition and promotes chemoresistance in pancreatic cancer cells, Oncogenesis 6 (2017) e352. 10.1038/oncsis.2017.54.

[25] C.R. Drifka, A.G. Loeffler, K. Mathewson, A. Keikhosravi, J.C. Eickhoff, Y. Liu, S.M. Weber, W. John Kao, K.W. Eliceiri, Highly aligned stromal collagen is a negative prognostic factor following pancreatic ductal adenocarcinoma resection, Oncotarget 7 (2016) 76197. 10.18632/ONCOTARGET.12772.

[26] A. Rubiano, D. Delitto, S. Han, M. Gerber, C. Galitz, J. Trevino, R.M. Thomas, S.J. Hughes, C.S. Simmons, Viscoelastic properties of human pancreatic tumors and in vitro constructs to mimic mechanical properties, Acta Biomater 67 (2018) 331–340. 10.1016/J.ACTBIO.2017.11.037.

[27] C.B. Raub, V. Suresh, T. Krasieva, J. Lyubovitsky, J.D. Mih, A.J. Putnam, B.J. Tromberg, S.C. George, Noninvasive assessment of collagen gel microstructure and mechanics using multiphoton microscopy, Biophys J 92 (2007) 2212–2222. 10.1529/biophysj.106.097998.

[28] C.C. Bell, A.C.A. Dankers, V.M. Lauschke, R. Sison-Young, R. Jenkins, C. Rowe, C.E. Goldring, K. Park, S.L. Regan, T. Walker, C. Schofield, A. Baze, A.J. Foster, D.P. Williams, A.W.M. van de Ven, F. Jacobs, J. van Houdt, T. Lähteenmäki, J. Snoeys, S. Juhila, L. Richert, M. Ingelman-Sundberg, Comparison of Hepatic 2D Sandwich Cultures and 3D Spheroids for Long-term Toxicity Applications: A Multicenter Study, Toxicological Sciences 162 (2018) 655–666. 10.1093/TOXSCI/KFX289.

[29] C. Jensen, Y. Teng, Is It Time to Start Transitioning From 2D to 3D Cell Culture?, Front Mol Biosci 7 (2020) 513823. 10.3389/FMOLB.2020.00033/REFERENCE.

[30] Y. Imamura, T. Mukohara, Y. Shimono, Y. Funakoshi, N. Chayahara, M. Toyoda, N. Kiyota, S. Takao, S. Kono, T. Nakatsura, H. Minami, Comparison of 2D- and 3D-culture models as drug-testing platforms in breast cancer, Oncol Rep 33 (2015) 1837–1843. 10.3892/OR.2015.3767/HTML.

[31] A. del M.B. Garnique, N.S. Parducci, L.B.L. de Miranda, B.O. de Almeida, L. Sanches, J.A. Machado-Neto, Two-Dimensional and Spheroid-Based Three-Dimensional Cell Culture Systems: Implications for Drug Discovery in Cancer, Drugs and Drug Candidates 2024, Vol. 3, Pages 391-409 3 (2024) 391–409. 10.3390/DDC3020024.

[32] M. Kanari, I. Jimenez Garcia, F.D. Steffen, L.A. Krattiger, C. Bataclan, W. Liu, B.R. Simona, B. Deplancke, O. Naveiras, M. Ehrbar, B. Bornhauser, J.-P. Bourquin, A three-dimensional ex vivo model recapitulates in vivo features and drug resistance phenotypes in childhood acute lymphoblastic leukemia, Leukemia 2025 (2025) 1–14. 10.1038/S41375-025-02739-8.

[33] H.A. Burris, M.J. Moore, J. Andersen, M.R. Green, M.L. Rothenberg, M.R. Modiano, M.C. Cripps, R.K. Portenoy, A.M. Storniolo, P. Tarassoff, R. Nelson, F.A. Dorr, C.D. Stephens, D.D. Von Hoff, Improvements in Survival and Clinical Benefit With Gemcitabine as First-Line Therapy for Patients With Advanced Pancreas Cancer: A Randomized Trial, Journal of Clinical Oncology 41 (2023) 5482–5492. 10.1200/JCO.22.02777.

[34] V. Heinemann, K. Grosshadern, Gemcitabine: Progress in the Treatment of Pancreatic Cancer, Oncology 60 (2000) 8–18. 10.1159/000055290.

[35] V. Heinemann, H. Wilke, H.G. Mergenthaler, M. Clemens, H. König, H.J. Illiger, M. Arning, A. Schalhorn, K. Possinger, U. Fink, Gemcitabine and cisplatin in the treatment of advanced or metastatic pancreatic cancer, Annals of Oncology 11 (2000) 1399–1403. 10.1023/A:1026595525977.

[36] H.M. Kocher, R. Georgescu, N. Kotriwala, C. Lawrence, G. Priyadarshini, C. Ackermann, C. Chelala, A. Imrali, C. Hughes, R. Roberts, D.K. Chang, J. Dixon-Hughes, P. Sasieni, P. Corrie, M.G. McNamara, D. Sarker, F.E.M. Froeling, A. Christie, R. Gillmore, K. Khan, D. Propper, Study protocol: multi-centre, randomised controlled clinical trial exploring stromal targeting in locally advanced pancreatic cancer; STARPAC2, BMC Cancer 25 (2025) 1–8. 10.1186/S12885-024-13333-Z/TABLES/1.

[37] G. Bi, J. Liang, Y. Bian, G. Shan, V. Besskaya, Q. Wang, C. Zhan, The immunomodulatory role of all-trans retinoic acid in tumor microenvironment, Clin Exp Med 23 (2023) 591–606. 10.1007/S10238-022-00860-X/FIGURES/2.

[38] H.M. Kocher, B. Basu, F.E.M. Froeling, D. Sarker, S. Slater, D. Carlin, N.M. deSouza, K.N. De Paepe, M.R. Goulart, C. Hughes, A. Imrali, R. Roberts, M. Pawula, R. Houghton, C. Lawrence, Y. Yogeswaran, K. Mousa, C. Coetzee, P. Sasieni, A. Prendergast, D.J. Propper, Phase I clinical trial repurposing all-trans retinoic acid as a stromal targeting agent for pancreatic cancer, Nat Commun 11 (2020) 1–9. 10.1038/S41467-020-18636-W;SUBJMETA.

[39] A. Chronopoulos, B. Robinson, M. Sarper, E. Cortes, V. Auernheimer, D. Lachowski, S. Attwood, R. Garciá, S. Ghassemi, B. Fabry, A. Del Rió Hernández, ATRA mechanically reprograms pancreatic stellate cells to suppress matrix remodelling and inhibit cancer cell invasion, Nat Commun 7 (2016) 1–12. 10.1038/NCOMMS12630;TECHMETA.

[40] M.R. Haque, C.R. Wessel, D.D. Leary, C. Wang, A. Bhushan, F. Bishehsari, Patient-derived pancreatic cancer-on-a-chip recapitulates the tumor microenvironment, Microsystems & Nanoengineering 2022 8:1 8 (2022) 1–13. 10.1038/s41378-022-00370-6.

[41] A. Armstrong, M.R. Haque, S. Mirbagheri, U. Barlass, D.Z. Gilbert, J. Amin, A. Singh, A. Naqib, F. Bishehsari, Multiplex patient-based drug response assay in pancreatic ductal adenocarcinoma, Biomedicines 9 (2021) 705. 10.3390/BIOMEDICINES9070705/S1.

[42] M. Huff, L. Romano, K. Jefimovs, Recent Advances in Reactive Ion Etching and Applications of High-Aspect-Ratio Microfabrication, Micromachines 2021, Vol. 12, Page 991 12 (2021) 991. 10.3390/MI12080991.

[43] M.S. Gerlt, N.F. Läubli, M. Manser, B.J. Nelson, J. Dual, Reduced etch lag and high aspect ratios by deep reactive ion etching (Drie), Micromachines (Basel) 12 (2021) 542. 10.3390/MI12050542/S1.

[44] Formlabs Design Specifications – SAIC Advanced Output Center, (n.d.). https://sites.saic.edu/aoc/formlabs-design-specifications/ (accessed December 10, 2024).

[45] A. Chronopoulos, B. Robinson, M. Sarper, E. Cortes, V. Auernheimer, D. Lachowski, S. Attwood, R. Garciá, S. Ghassemi, B. Fabry, A. Del Rió Hernández, ATRA mechanically reprograms pancreatic stellate cells to suppress matrix remodelling and inhibit cancer cell invasion, Nature Communications 2016 7:1 7 (2016) 1–12. 10.1038/ncomms12630.

[46] GitHub - gnea/grbl-Mega: An open source, embedded, high performance g-code-parser and CNC milling controller written in optimized C that will run on an Arduino Mega2560, (n.d.). https://github.com/gnea/grbl-Mega (accessed March 21, 2024).

[47] J. Rajagopalan, M.T.A. Saif, Fabrication of freestanding 1-D PDMS microstructures using capillary micromolding, Journal of Microelectromechanical Systems 22 (2013) 992–994. 10.1109/JMEMS.2013.2262605.

[48] Corning Incorporated, Certificate of Analysis for Corning Collagen I High concentration (HC), Rat Tail, (n.d.). https://certs-ecatalog.corning.com/life-sciences/certs/354249_9343002.pdf (accessed July 5, 2020).

[49] D. Sharma, D. Adnan, M.K. Abdel-Reheem, R.C. Anafi, D.D. Leary, F. Bishehsari, Circadian transcriptome of pancreatic adenocarcinoma unravels chronotherapeutic targets, JCI Insight 9 (2024). 10.1172/JCI.INSIGHT.177697.

